# The augmentation of retinogeniculate communication during thalamic burst mode

**DOI:** 10.1101/427674

**Authors:** Henry Alitto, Daniel L. Rathbun, Jessica J. Vandeleest, Prescott C. Alexander, W. Martin Usrey

**Author notes:** <u>Correspondence</u>: Henry J. Alitto, PhD, Center for Neuroscience, University of California, Davis, 1544 Newton Court, Davis, CA 95618, USA.

## Abstract

Retinal signals are transmitted to cortex via neurons in the lateral geniculate nucleus (LGN), where they are processed in burst or tonic response mode. Burst mode occurs when LGN neurons are sufficiently hyperpolarized for T-Type Ca^2+^ channels to de-inactivate, allowing them to open in response to a depolarization which can trigger a high-frequency sequence of Na+-based spikes (i.e. burst). In contrast, T-type channels are inactivated during tonic mode and do not contribute to spiking. Although burst mode is commonly associated with sleep and the disruption of retinogeniculate communication, bursts can also be triggered by visual stimulation, thereby transforming the retinal signals relayed to the cortex.

To determine how burst mode affects retinogeniculate communication, we made recordings from monosynaptically connected retinal ganglion cells and LGN neurons in the cat during visual stimulation. Our results reveal a robust augmentation of retinal signals within the LGN during burst mode. Specifically, retinal spikes were more effective and often triggered multiple LGN spikes during periods likely to have increased T-Type Ca^2+^ activity. Consistent with the biophysical properties of T-Type Ca^2+^ channels, analysis revealed that effect magnitude was correlated with the duration of the preceding thalamic interspike interval and occurred even in the absence of classically defined bursts. Importantly, the augmentation of geniculate responses to retinal input was not associated with a degradation of visual signals. Together, these results indicate a graded nature of response mode and suggest that, under certain conditions, bursts facilitate the transmission of visual information to the cortex by amplifying retinal signals.

**Significance:** The thalamus is the gateway for retinal information traveling to the cortex. The lateral geniculate nucleus (LGN), like all thalamic nuclei, has two classically defined categories of spikes—tonic and burst—that differ in their underlying cellular mechanisms. Here we compare retinogeniculate communication during burst and tonic response modes. Our results show that retinogeniculate communication is enhanced during burst mode and visually evoked thalamic bursts, thereby augmenting retinal signals transmitted to cortex. Further, our results demonstrate that the influence of burst mode on retinogeniculate communication is graded and can be measured even in the absence of classically defined thalamic bursts.

## Introduction

The lateral geniculate nucleus (LGN) of the dorsal thalamus is the primary source of visual signals sent to primary visual cortex (V1), receiving monosynaptic input from retinal ganglion cells (RGCs) and projecting directly to cortical target neurons. Despite being labeled a relay nucleus, the LGN serves to transform retinal signals in several significant and dynamic ways (Dan et al., 1996; Usrey et al., 1998, Martinez et al., 2014; Fisher et al., 2017; Alitto et al., 2018), including changes in the temporal domain that accompany tonic and burst activity modes (reviewed in Sherman and Guillery, 2009; Usrey and Alitto, 2015). During tonic mode, LGN neurons respond to excitatory input with regularly spaced action potentials, the rate of which is proportional to the strength of the stimulus (Llinas and Jahnsen, 1982, Huguenard and McCormick, 1992). By contrast, LGN spike trains during burst mode are irregular, include tight clusters of spikes known as "bursts”, and firing rate becomes uncoupled from stimulus strength. Although geniculate bursts are generally associated with periods of low arousal and sleep, when LGN neurons are thought to be dissociated from the periphery, they can also occur during sensory processing and have been shown to be particularly effective in evoking cortical responses (Swadlow and Gusev, 2001; Weyand et al., 2001; Lesica and Stanley, 2004; Alitto and Usrey, 2005; Denning and Reinagel, 2005; Bezdudnaya et al., 2006; Alitto and Usrey, 2011; Bereshpolova et al., 2011). Determining how burst mode affects retinogeniculate communication is therefore important for understanding the transmission of visual information to the cortex.

Across thalamic nuclei the transition from tonic to burst mode depends on a common mechanism, the de-inactivation of T-type Ca^2+^ channels (or T-channels) that occurs when neurons are sufficiently hyperpolarized for a sufficient duration of time (Llinas and Jahsen, 1982; Huguenard and McCormick, 1992; Wei et al., 2011). When this occurs, depolarizing stimuli can activate T-channels to generate a suprathreshold Ca^2+^ potential (T-potential), which can then trigger a short train of high-frequency, Na+-based action potentials. It is important to note that the magnitude of the T-potential and subsequently the number of spikes it triggers depends on the percentage of T-channels in the de-inactivated versus the inactivated state which, in turn, depends on the depth and duration of the preceding hyperpolarization (Deschenes et al., 1984; Destexhe et al., 2002; Hong et al., 2014).

Here, we explore the influence of thalamic response mode on retinogeniculate communication by performing simultaneous extracellular recordings of monosynaptically connected pairs of RGCs and LGN neurons in the anesthetized cat. Although the occurrence of T-potentials is best determined with intracellular recording methods, past work has shown that bursts can be identified by applying a previously established set of statistical criteria to extracellular records of LGN spike trains (Lu et al., 1992; see Materials and Methods). Using these criteria, we calculated retinal efficacy (percentage of RGC spikes that triggered LGN cell spikes) and retinal contribution (percentage of LGN spikes evoked by a simultaneously recorded RGC) during tonic and burst response modes. Our results reveal a fundamental change in retinogeniculate communication during burst mode and suggest an augmentation of visual signals by T-potentials. We found that individual retinal spikes arriving during epochs supportive of T-channel activity were more effective in evoking LGN responses and often triggered multiple spikes. Further, there was a decrease in the percentage of LGN responses directly triggered by retinal spikes during thalamic bursts; however, this was not associated with a degradation of visual signals within the LGN. Consistent with the biophysical properties of T-channels, the modulation of retinogeniculate communication was proportional to the duration of the preceding interspike interval of the LGN neuron and was evident even in the absence of classically defined thalamic bursts. These results reveal how retinal signals are transformed by the transition between tonic and burst modes and, importantly, suggest that the influence of thalamic response mode on retinogeniculate communication is a continuous process.

## Materials and Methods

### Animal preparation

Sixteen adult cats were used for this study. All experimental procedures were conducted with the consent of the Animal Care and Use Committee at the University of California, Davis and followed NIH guidelines. Some of the data analyzed in this study contributed to previous unrelated studies on the retinogeniculate pathway (Rathbun et al., 2010; Rathbun et al., 2016; Usrey et al., 1998,1999).

Surgical procedures were preformed while animals were anesthetized. Surgical anesthesia was induced with ketamine (10 mg/kg, IM) and maintained with thiopental sodium (20 mg/kg, IV, supplemented as needed). A tracheotomy was performed and animals were placed in a stereotaxic apparatus where they were mechanically ventilated. EEG, EKG, CO^2^ and temperature were monitored throughout the experiment. A scalp incision was made and wound edges were infused with lidocaine. A craniotomy was made over the LGN, the dura was removed, and the craniotomy was filled with agarose to protect the underlying brain. Eyes were adhered to metal posts, fitted with contact lenses, and focused on a tangent screen located 172 cm in front of the animal. Phenylephrine (10%) was administered to retract the nictitating membranes and flurbiprofen sodium drops were administered (1.5 mg/hr) to prevent miosis. The positions of area centralis and the optic disk were mapped by back-projecting the retinal vasculature of each eye onto a tangent screen. After the completion of surgical procedures, maintenance anesthesia (thiopental sodium (2-3 mg/kg/hr, IV) was administered for the remaining duration of the experiment. Supplemental thiopental was given and the rate of infusion was increased if physiological monitoring indicated a decrease in the level of anesthesia. Once a steady plane of maintenance anesthesia was established, animals were paralyzed with vecuronium bromide (0.2 mg/Kg/hr, IV). Animals were euthanized with Euthasol (100 mg/kg; Virbac Animal Health, Fort Worth, Texas) at the conclusion of each experiment.

### Electrophysiological recording and visual stimuli

Simultaneous extracellular recordings were made from LGN cells in layers A and A1 and retinal ganglion cells. For thalamic recordings, the LGN was first located using single, parylene-coated tungsten electrodes (AM Systems, Everett, WA). After the preferred retinotopic position was located in the LGN, a 7-channel multielectrode array (Thomas Recording, Marburg, Germany) was positioned into the LGN. Retinal ganglion cells were recorded from using a tungsten-coated microelectrode inserted into the eye through an intraocular guide tube and maneuvered via a custom-made manipulator. Neural responses were amplified, filtered and recorded to a computer equipped with a Power 1401 data acquisition interface and the Spike 2 software package (Cambridge Electronic Design, Cambridge, UK). Spike isolation was based upon waveform analysis (parameters established independently for each cell) and the presence of a refractory period as indicated in the autocorrelogram (Usrey et al., 2000, 2003).

Visual stimuli were generated using a VSG2/5 visual stimulus generator (Cambridge Research Systems, Rochester, England) and presented on a gamma-calibrated Sony monitor running at 140Hz. The mean luminance of the monitor was 38 candelas/m^2^. Visual responses of LGN neurons and RGCs were mapped and characterized using drifting sine-wave gratings and white-noise stimuli. The white-noise stimulus consisted of a 16×16 grid of black and white squares. Each square was independently modulated in time according to an m-sequence of length 2^15^-1 (Sutter, 1992; Reid et al., 1997). Individual squares in the stimulus were updated with each monitor frame for 2^15^ -1 frames (~4 minutes). Approximately 4-16 squares of the stimulus overlapped each neuron’s receptive field center. Drifting sine-wave grating stimuli (4 Hz, 100% contrast) were presented at the preferred spatial frequency for the recorded cells.

### Cross-correlation analysis

Cross-correlograms between retinal and geniculate spike trains were made to assess connectivity between pairs of cells (Figure 1). Cross-correlograms were calculated by generating histograms of LGN spikes relative to each retinal spike (Figure 1A) and retinal spikes relative to each LGN spike (Figure 1B). Peaks indicative of monosynaptic connectivity were narrow (< 1.5 ms, full width at half height), short-latency (< 5 ms), and exceeded 5× the standard deviation of the baseline (Cleland et al., 1971; Usrey et al. 1998). For quantitative analysis, bins contributing to the peak were identified using a bin size of 0.5 ms. The peak bin was first identified and all neighboring bins greater than 3 standard deviations above the baseline mean were considered part of the peak, where the baseline consisted of bins ranging from 30 to 50 ms on either side of the peak bin.

**Figure 1.**
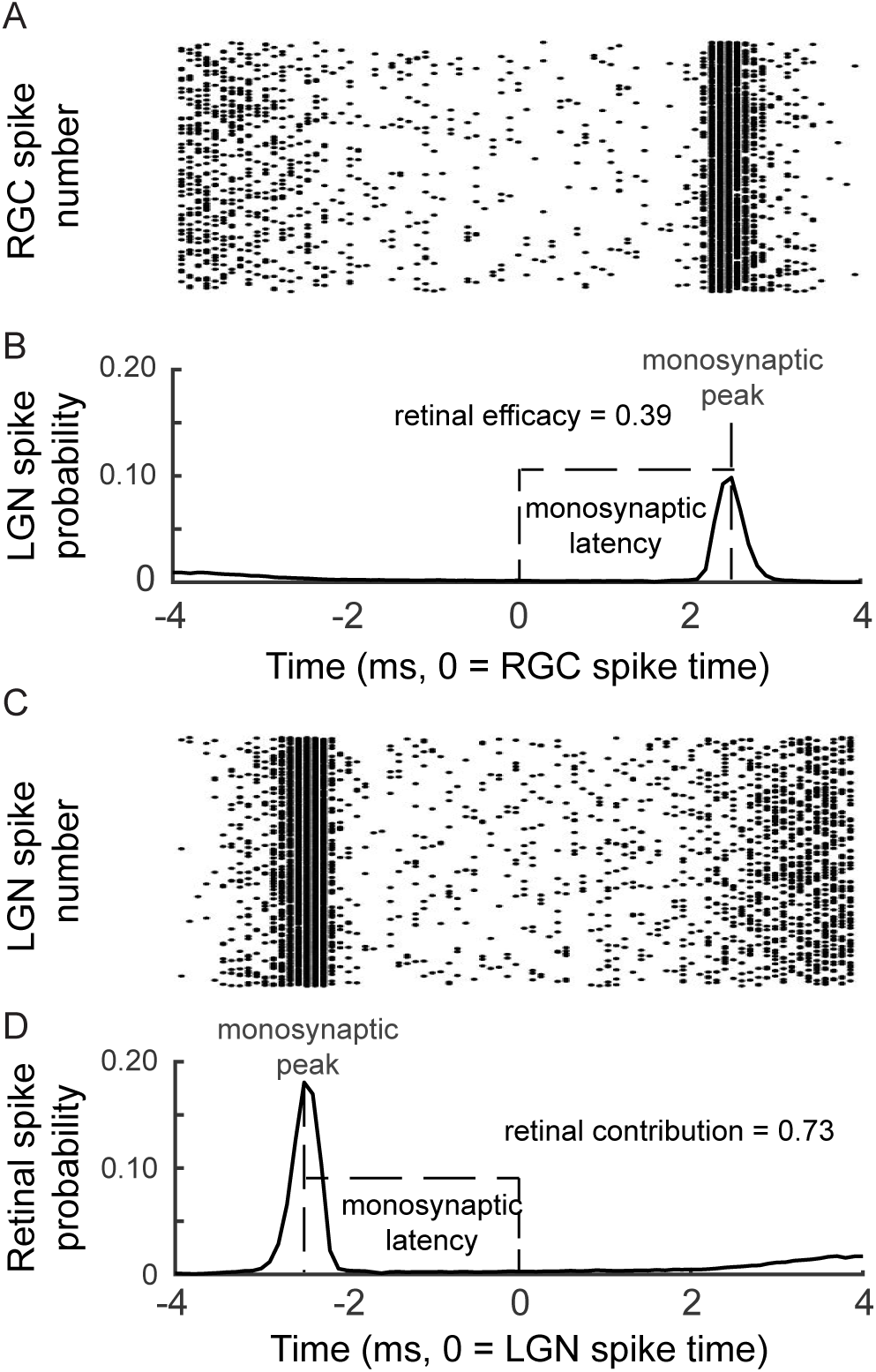
Cross-correlation analysis to identify monosynaptic connections between retinal ganglion cells and LGN neurons. ***A***, A raster plot showing the timing of action potentials of an LGN neuron relative to the action potentials of a simultaneously recorded RGC (time = 0). ***B***, A clear, narrow, short-latency peak can be seen in this example cross-correlogram, indicating a monosynaptic connection between the two neurons. For this example, retinal efficacy was 0.39. ***C, D***, Similar plots showing the timing of action potentials of the RGC relative to those of the LGN neuron (time = 0). For this example, retinal spike contribution was 0.73.

### Retinal spike contribution and efficacy

Cross-correlation analysis was used to assess connectivity between cell pairs as well as strength of connection. The monosynaptic peak in a cross correlogram was used to calculate two measures of correlation strength, efficacy (Figure 1B) and contribution (Figure 1D; Cleland et al., 1971; Usrey et al. 1998). Efficacy is the number of spikes in the monosynaptic peak divided by the total number of retinal spikes; contribution is the number of spikes in the peak divided by the total number of LGN spikes. To the extent that peaks were caused by monosynaptic connections, efficacy and contribution have very simple interpretations. Efficacy represents the fraction of the retinal spikes that evoked geniculate spikes, and contribution represents the fraction of the geniculate spikes that were caused by a spike from the RGC. Given that LGN neurons receive convergent input from 2-6 RGCs, it is worth noting that this measure of retinal contribution quantifies the influence of the simultaneously recorded RGC on the spiking behavior of the LGN neuron and not the combined influence of all of the RGCs that provide convergent input to the LGN neuron.

When applicable we calculated the expected retinal spike efficacy for each recorded cell pair as follows (Alitto et al., 2018). First, we calculated the average spike efficacy across a range of interspike intervals (ISIs), estimated independently for responses driven by drifting gratings and white noise. We then modeled the expected spike efficacy by assigning each retinal spike the efficacy value calculated for the corresponding ISI. Thus, the spike efficacy became the value expected if retinogeniculate transmission did not systematically depend upon a given independent variable.

### Identification of LGN bursts and tonic spikes

We used two well-established criteria to identify bursts in the spike trains of LGN neurons (Lu et al., 1992; Swadlow and Gusev, 2001; Weyand et al., 2001; Lesica and Stanley, 2004; Alitto and Usrey, 2005; Denning and Reinagel, 2005; Bezdudnaya et al., 2006; Alitto and Usrey, 2011; Bereshpolova et al., 2011). These criteria were: (1) an interspike interval (ISI) >100 ms that preceded the first spike in a sequence and (2) one or more subsequent spikes that followed with ISIs < 4 ms (Figure 2A). Past studies applying these criteria to intracellular recordings show that events defined as bursts co-occur with T-channel plateau potentials (Lu et al., 1992). For this study, the first spike in the burst is referred to as the cardinal spike, and each additional spike is referred to by its ordinal position (secondary, tertiary, etc.).

**Figure 2.**
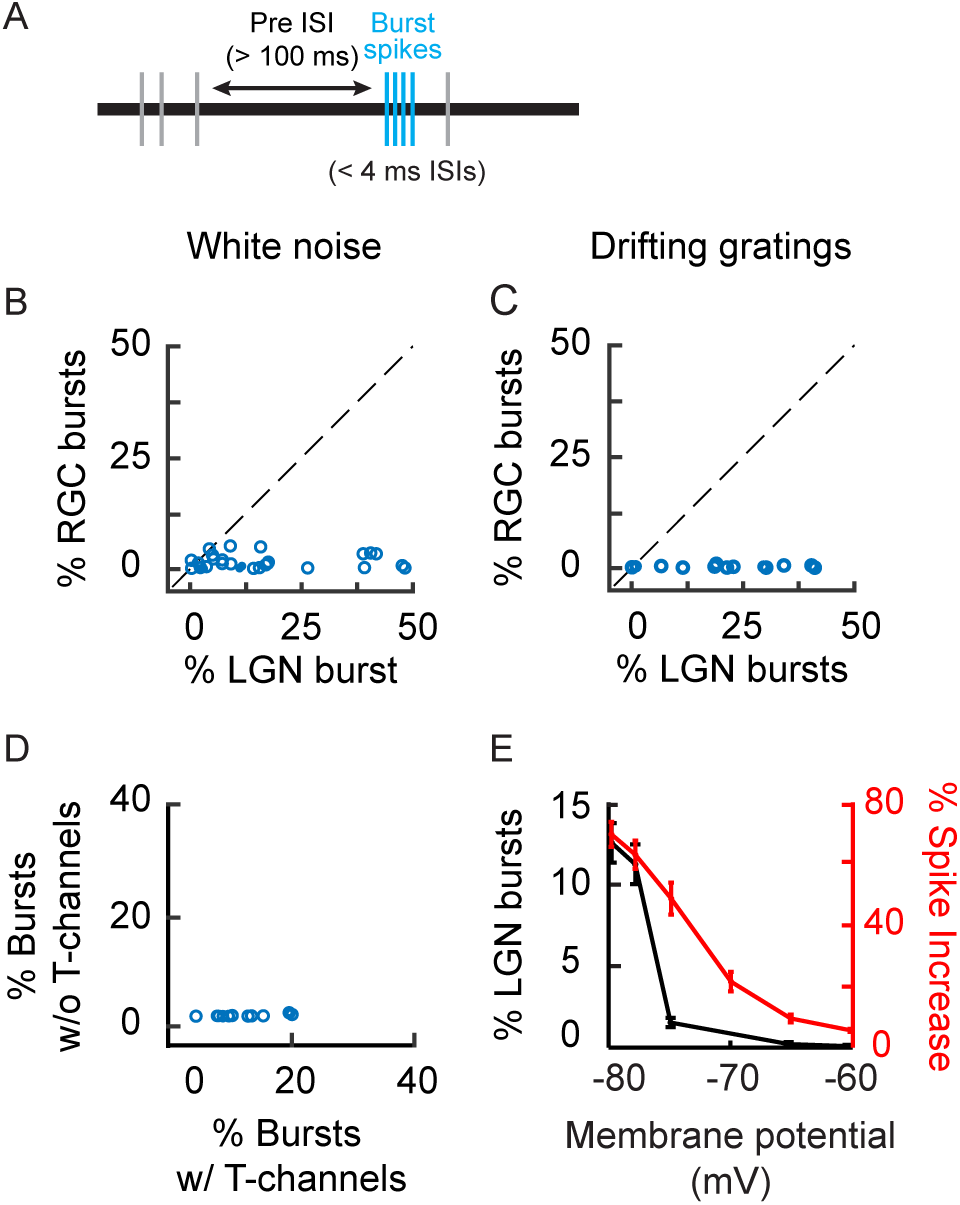
Comparison of burst frequency in the retina and LGN. ***A***, Bursts (blue tick marks) were identified by applying the following criteria to extracellular recordings: (1) the first spike was preceded by an interspike interval (ISI) >100 ms (horizontal arrow) and (2) subsequent spikes followed with ISIs < 4 ms. ***B, C***, Scatterplot showing the percentage of RGC and LGN cell spikes that were identified as part of a burst, during white-noise (***B***) and drifting grating (***C***) stimulation. ***D***, Scatterplot showing the percentage of simulated LGN spikes that were identified as part of a burst when a leaky integrate-and-fire mode either included or did not included T-channels. ***E***, Line graph showing the influence of membrane potential on the percentage of LGN spikes that were identified as part of a burst when the simulation included T-channels (left y-axis, black line) and the increase in simulated LGN spike count due to the addition of T-channels to the model (right y-axis, red line). Error bars = standard error.

### Simulation of T-type Ca^+2^ channels

Given the critical role that T-type Ca^+2^ channels play in the generation of thalamic bursts and that the biophysical properties of these channels have been extensively characterized, we simulated the interaction of T-Type Ca^+2^ channels and synaptic EPSPs. We used a leaky-integrate and fire model neuron and a series of previously published equations that quantify the voltage and time dependence of both the de-inactivation and inactivation of T-Channels (Huguenard and McCormick, 1992). There is at least one other variation of this series of equations (Wang et al., 1991); however, the two versions produce equivalent results within the scope of the current study. The membrane potential of the model neuron was simulated by:

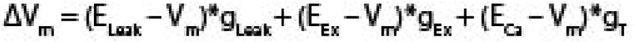

Here, E_Leak_, E_Ex_, and E_Ca_ are the reversal potentials for the leak current, excitatory input, and T-Channels, respectively. g_Leak_, g_Ex_, and g_T_ are the conductance values for the leak current, the excitatory inputs, and the T-channels. Synaptic inhibition was not necessary to produce thalamic bursts, so they were not included in this simulation. Excitatory input was simulated using the retinal spike trains recorded *in vivo*. When the Vm exceeded -35 mV an action potential was recorded and Vm was reset to -60 mV. Maximum g_Ex_ was selected to generate biologically reasonable firing rates and retinal spike-efficacy curves with a -60 mV resting membrane potential. g_T_ was controlled by the following voltage and time dependent equations:

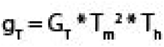

Here, G_T_ is the maximum T-channel conductance, T_m_ is the activation gate and T_h_ is the inactivation gates for the T-channels.

The activation states of T_m_ and T_h_ were determined by the following equations:

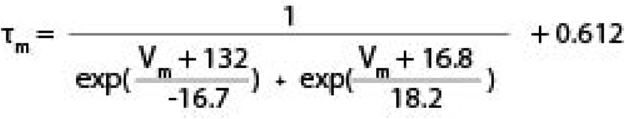

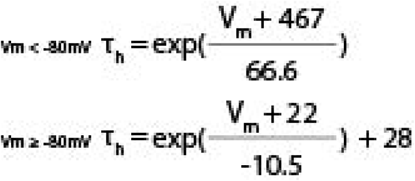

Here τ_m_ and τ_h_ are the time constants for the activation and inactivation gates, respectively.

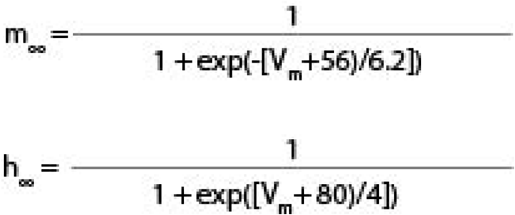

Here, m∞ and h∞ are the steady state activation levels for the activation and inactivation gates, respectively. For more details on simulating T-channels see Huguenard and McCormick, 1992, Smith et al., 2000, and Destexhe and Sejnowski, 2001.

### Spatiotemporal receptive field maps

Spatiotemporal receptive fields (STRFs) were calculated from LGN spike trains evoked during the presentation of a binary white-noise stimulus (16 × 16 grid of black and white squares). Each square was temporally modulated according to a 2^15^-1 length m-sequence <u>(Reid and Shapley, 1992; Sutter, 1992; Reid et al., 1997)</u>. The stimulus was updated either every frame (7.1 ms) or fourth frame of the display (28.4 ms), and the entire sequence (~4 or 16 min) was typically repeated, up to 10x. To determine if LGN burst spikes were driven by visual stimulation, LGN STRFS were calculated using either the full spike trains (all spikes) or spike-count matched subsets of data (e.g., only cardinal burst spikes). Spike-count matching was done on a cell-by-cell basis by determining which subset had the least spikes and then randomly subsampling the other subsets to have the same total. This was done so that the signal-to-noise ratios (STN) were comparable within a given cell. The signal was estimated as the amplitude of the 2D Gaussian fit (Matlab function fmincon) to the frame of the STRF containing the peak pixel. The Gaussian receptive field estimate is described by the following equation: G_ij_ = K × exp[-(x_i_ – x_0_)^2^/2 × σ^2^] × exp[-(y_i_ – y_0_)^2^/2 × σ2], where K is the amplitude, x_0_ and y_0_ are the coordinates of the center of the receptive field, and σ is the standard deviation. Noise was estimated as the mean value for three frames centered at t = +100ms.

### Experimental design and statistical analysis

To quantify the relationship between retinogeniculate communication and thalamic response mode we used generalized linear mixed effect models (GLME, Matlab function fitglme Raudenbush and Bryk, 2001) using a Laplace fit method. This is done to take full advantage of the number of data points collected (e.g., hundreds of thousands of retinal and thalamic spikes) while accounting for differences between cells. The general form of a GLME is:

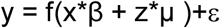

Here y is the outcome being modeled, x is matrix of fixed effects variables, β is a vector of fixed effects coefficients, z is a matrix of random effects variables, μ is a vector of random effects coefficients, ε is the residual error, f is the link function. β_0_ is the y-intercept, while β_variable name_ is the coefficient of a specific variable (e.g., β_ISI_). When analyzing the percentage of high-frequency spikes, these values were modeled using the identify function and as arising from a normal distribution. In this case, the β coefficients represent the linear slope between the predicted outcome and the fixed effect variable. When analyzing the percentage of spikes per burst, these values were modeled using a log link function and as arising from a Poisson process. When analyzing retinal contribution and retinal spike efficacy, these values were modeled using a logit link function and as arising from a Bernoulli process (0s and 1s). For example, retinal spikes were assigned values of 1s and 0s based on whether they triggered an LGN action potential (1) or did not (0), as described above. In this case, the β coefficients represent the influence of the fixed effect variable on the log of the odds ratio of the predicted outcome. For each GLME model, cell identity was set as a random effect to account for differences between cells. For illustrative purposes, data was binned and normalized (e.g., Figure 6). Normalization was performed such that the average value (efficacy or contribution) was set to 1.0. It is important to note that these transformations were done to represent effects graphically that are difficult to directly represent based on Bernoulli variables; however, the GLMEs models were fit to the raw values that were neither binned nor normalized.

When simpler statistical analyses were sufficient to compare two distributions, we first tested the normality of the distributions using Lilliefors modification of the Kolmogorov-Smirnov test. If it was determined that both distributions were not significantly different from normal distributions, then a t-test was used to compare the means of the two samples, otherwise a Wilcoxon rank sum test or a sign test was used. X and Y cells were classified based on the latency of the monosynaptic peak (Usrey et al., 1999). Using this measure, of the 29 cell pairs examined in this study, 7 were X cell pairs and 22 were Y cell pairs. Results did not differ for these cell groups; thus the 29 cell pairs were treated as a single group for the statistical analyses presented. It should be noted that small differences between X and Y cells may have gone undetected because of the small sample sizes inherent to studying monosynaptic connections *in vivo*.

## Results

To quantify retinogeniculate communication during tonic and burst activity modes in the LGN, we made simultaneous recordings of synaptically connected RGCs and LGN neurons in the anesthetized cat. Retinal and geniculate neurons were excited with white-noise stimuli (n=29 cell pairs) and/or drifting sinusoidal gratings (n=15 cell pairs; see Materials and Methods). As will be expounded upon in the discussion section, these stimuli were chosen because of how their spatiotemporal profiles might differentially interact with geniculate response mode. Geniculate bursts were identified using established criteria for extracellularly recorded spikes (Lo et al., 1991). Specifically, a burst was defined as a sequence of spikes that met two criteria (Figure 2A): (1) the first spike in the sequence followed an ISI > 100 ms, and (2) one or more subsequent spikes followed with ISIs < 4 ms. Across 29 monosynaptically connected pairs of RGCs and LGN neurons, we recorded 1,394,029 retinal spikes and 530,428 LGN spikes, including 54,482 geniculate bursts (2 or more spikes). As expected, burst frequency was significantly greater for LGN neurons than for RGCs (Figure 2B-C; during white-noise stimulation: RGC = 1.5% +/-0.3%; LGN = 16.1% +/-2.8%, p < 10^-5^; during drifting grating stimulation: RGC =0.24% +/-0.6%, LGN = 26.2% +/-4.7%, p < 10^-5^).

### Simulating thalamic bursts involving T-potentials

Given that the biophysical properties of T-channels are well characterized, simulations can be used to illustrate how T-channels and retinal spikes are predicted to interact and transform retinogeniculate communication (see Materials and Methods). In particular, leaky integrate and fire neuronal models generate bursts with the simple addition of T-channels based on published equations (Figure 2D, Percent Burst with T-channels: 11.5±0.3%, percent burst without T-channels: 0.1±0.05%, p<10^-5^, see Methods; Huguenard and McCormick, 1992). Similarly, the addition of T-channels increased the number of geniculate spikes evoked from the same excitatory input (Figure 2E, blue line and axis). Interestingly, the increase in geniculate spike count remained elevated at higher resting membrane potentials where the percentage of burst spikes was greatly reduced (Figure 2E, black line and axis). This suggests that the influence of T-channels on geniculate activity can be measured even in the absence of classically defined bursts.

We hypothesized that visually evoked T-potentials will augment the transmission of visual signals through the LGN because of the summation of retinal EPSPs with T-potentials. Specifically, T-potentials are predicted to increase the ability of retinal spikes to trigger geniculate spikes as well as cause single retinal spikes to trigger multiple LGN spikes. Further, given the voltage and time dependence of the de-inactivation of T-channels, the influence of T-potentials on retinogeniculate communication should be dynamically regulated by the depth and duration of the preceding membrane hyperpolarization (Figure 3).

**Figure 3.**
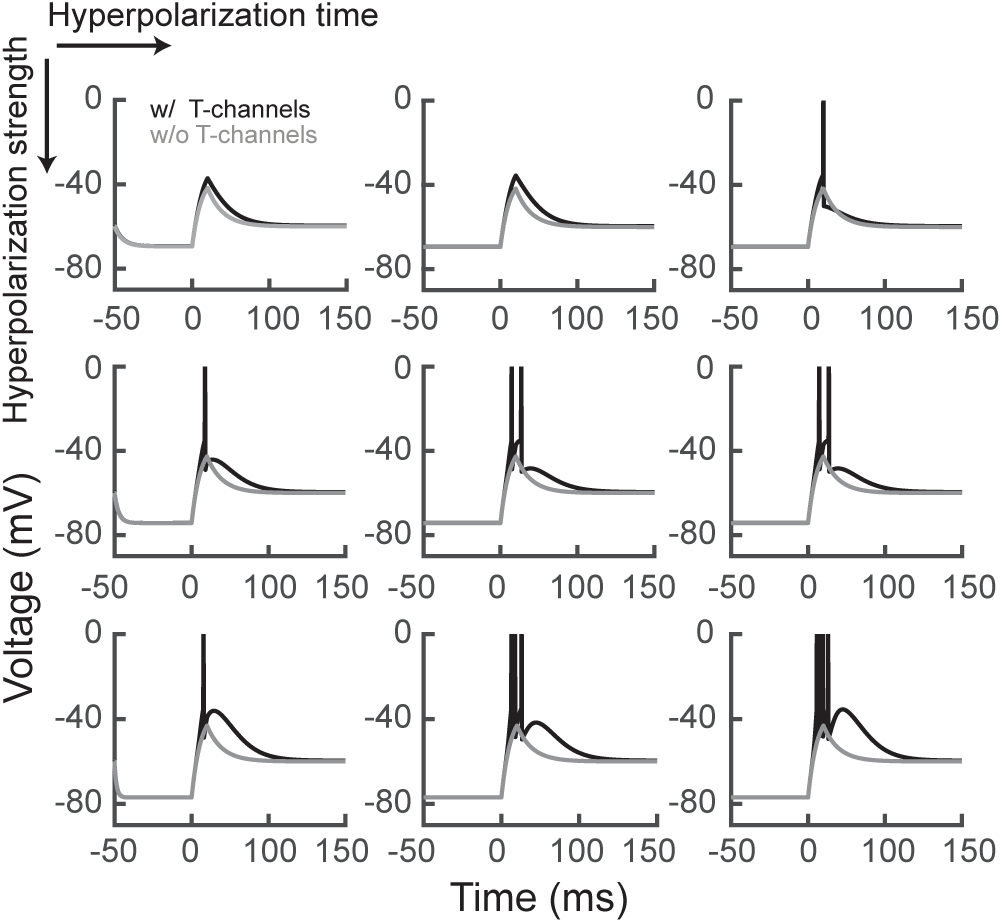
Leaky integrate-and-fire simulation of geniculate bursts. Simulation of T-potentials using a standard integrate-and-fire neural model. Using previously published equations (see Materials and Methods) we simulated the influence of increasing the amplitude (progressively stronger by row) and duration (progressively longer by column) of a hyperpolarization on T-channel activation to response to depolarization. Black lines = model with T-channels, gray lines = model without T-channels.

Although membrane hyperpolarization cannot be directly measured from extracellular recordings, it is likely that its influence on T-channel activity is correlated with the length of the LGN cell’s preceding ISI—the longer the ISI, the greater the probability of T-channel de-inactivation. To test this idea, we compared the probability that LGN cells generate high-frequency spikes (ISIs less than 4 ms) as a function of the LGN cells’ preceding ISI. Results show that percentage of high-frequency spikes from LGN cells increased dramatically as the preceding ISI increased beyond 50 ms (Figure 4A and B, white noise, β_ISI_ = 0.59±0.05, p < 10^-5^, dist. = normal; drifting grating, β_ISI_ = 0.59±0.15, p < 0.0005, dist. = normal). This effect was not seen for RGCs. Similarly, the number of spikes per burst was also directly dependent upon the preceding ISI (Figure 4C and D; white noise: β_ISI_ = 0.65±0.1, p < 10^-5^, dist. = Poisson; drifting grating, β_ISI_ = 0.29±0.14, p < 0.053, dist. = normal).

**Figure 4.**
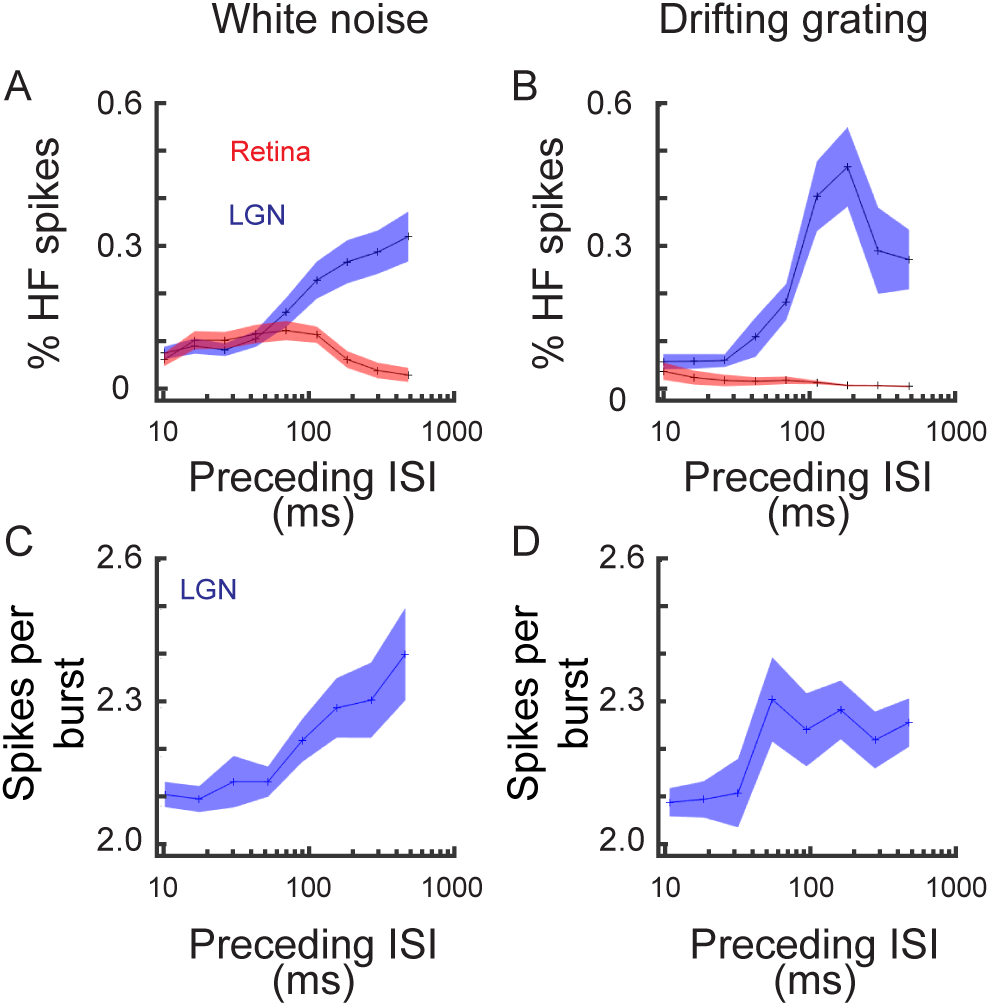
The influence of preceding ISI on high-frequency spiking in the retina and LGN. ***A, B***, Line plots showing the influence of preceding ISI on the percentage of high-frequency spikes (red line = RGC, blue line = LGN) during white-noise (***A***) and drifting grating (***B***) stimulation. High-frequency spikes are defined as two or more consecutive spikes with ISIs <4 ms. Shaded area = stand error. ***C, D***, Line plots showing the influence of preceding LGN ISI on the number of spikes per burst.

### Visually triggered geniculate bursts

The de-inactivation of T-channels that is fundamental to thalamic bursts can occur via multiple mechanisms. During sleep and deep anesthesia, when thalamic neurons typically fire in burst mode, the de-inactivating hyperpolarization is not associated with visual stimulation but rather involves intrinsic corticothalamic oscillations. Under these conditions, intrinsically generated bursts decouple the retina from the LGN and therefore do not convey visual information to the cortex. However, as we and others have shown previously, the hyperpolarization needed to de-inactivate T-channels can also result from visual stimulation (Alitto et al., 2005; Denning and Reinagel, 2005; Ortuño et al., 2014). Under these conditions, bursts do not degrade visual signals, but instead relay retinal/visual information to the cortex. Given these very different mechanisms for burst production and the implications each mechanism would have on the interpretation of our data, we examined the spike trains of the cells in this study to determine whether or not the bursts conveyed visual information. To do so, we calculated space-time receptive fields from LGN responses to the white-noise stimulus using only burst spikes and compared these response maps to those computed using a spike-count matched subset of tonic spikes. As shown in Figure 5 (A and B), burst and tonic maps had similar signal-to-noise ratios indicating that the burst spikes were evoked by visual stimulation (tonic spikes: 9.5±1.5, burst spikes: 10.6±1.3, p=0.54). Similarly, burst spikes recorded during visual stimulation with drifting gratings were tightly phase locked to the stimulus (Figure 5C and D; tonic spikes circular variance: 0.12±0.02, burst spikes: 0.03±0.01, p=0.001).

**Figure 5.**
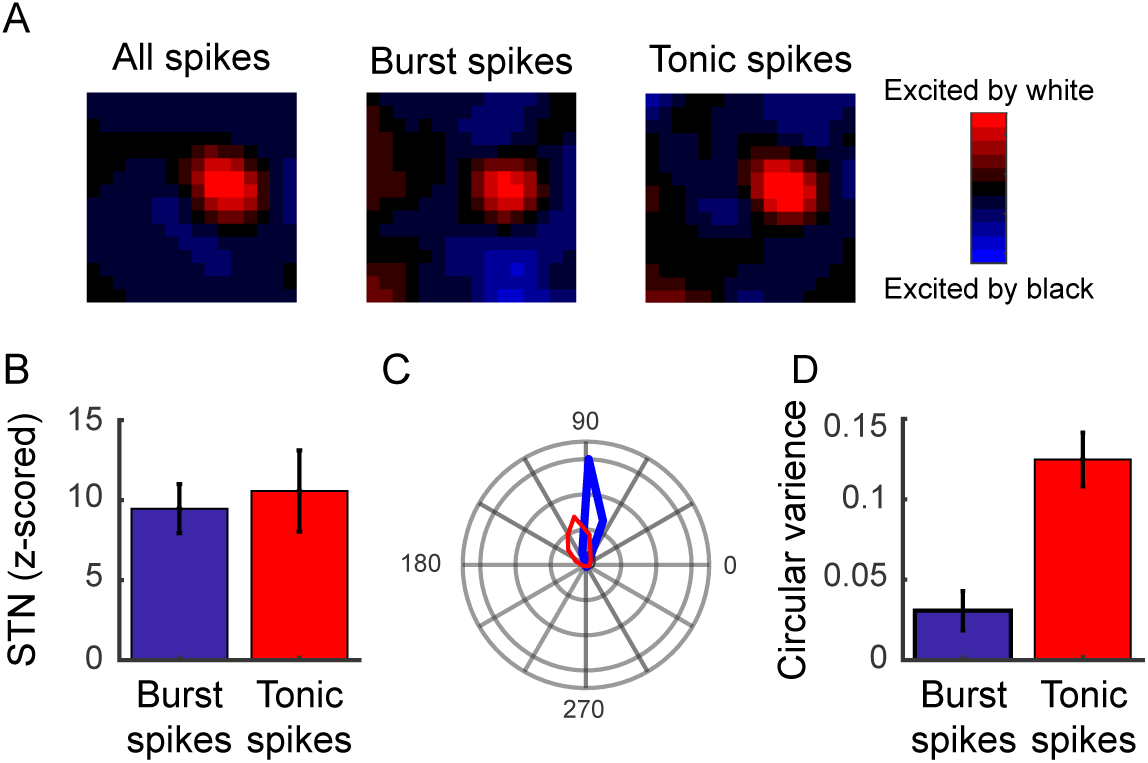
Geniculate bursts are evoked by visual stimulation. ***A***, Spatiotemporal receptive field (STRF) maps from a representative LGN neuron calculated using specific subsets of spike-count matched geniculate spikes: all spikes (left), burst spikes (middle), and tonic spikes (right). ***B***, Bar graph showing sample mean signal-to-noise ratios (STN) for tonic and burst STRFs. ***C***, Polar plot illustrating the phase locking of LGN tonic (red line) and burst (blue line) spikes during visual stimulation with drifting gratings. ***D***, Bar graph showing circular variance for tonic and burst spikes during visual stimulation with drifting gratings. Low circular variance values indicate that the spikes were phase locked to the visual stimulus, while a value of 1 indicates that the spikes occurred equally across all phases. Error bars = standard error.

### Augmentation of retinal signals during visually triggered geniculate bursts

To test the hypothesis that visually evoked geniculate bursts are associated with an amplification of the retinal signal within the LGN, we measured retinal spike efficacy as a function of time since the most recent LGN spike. Using the assumption that the probability of T-channel de-inactivation increases as the LGN ISI increases in duration (see above), we calculated retinal spike efficacy as a function of the "ongoing” LGN ISI (Figure 6A). For example, if a retinal spike occurred 10 ms after the most recent LGN spike, the ongoing LGN ISI is 10 ms, regardless of the timing of either the previous retinal spike or the next thalamic spike. If T-channels de-inactivate during relatively long LGN ISIs, then retinal spikes that occurred during such ongoing LGN ISIs are predicted to trigger T-potentials and thus have an enhanced ability to evoke a geniculate response.

**Figure 6.**
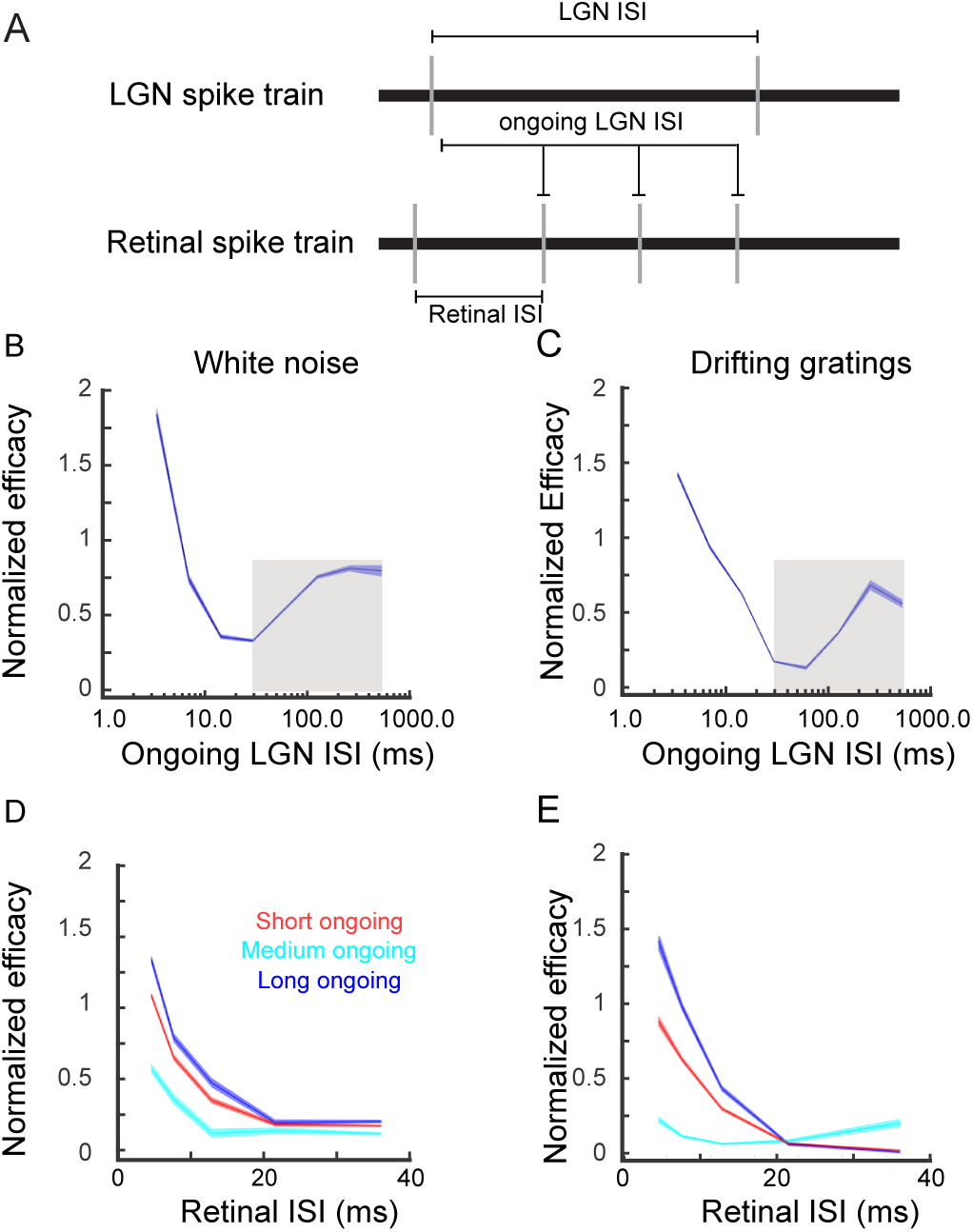
Retinal spike efficacy is influenced by ongoing LGN ISI. ***A***, Ongoing LGN ISI is defined as the time since the most recent LGN spike at the occurrence of a RGC spike. This is in contrast to retinal ISI, the interval between two consecutive RGC spikes, and LGN ISI, the interval between two consecutive LGN spikes. ***B***, ***C***, Line plots showing the influence of ongoing LGN ISI on retinal spike efficacy, during white-noise (***B***) and drifting grating stimulation (***C***). The shaded areas around the line indicate standard error. The gray boxes indicate the range of ISI values used for the GLME model (see Materials and Methods). ***D, E***, Line plots showing the influence of ongoing LGN ISI on retinal spike efficacy (red = ongoing LGN ISI < 30ms, light blue line = ongoing LGN ISI > 30ms and < 100ms, dark blue line = ongoing LGN ISI > 100ms).

Unsurprisingly, retinal spike efficacy was greatest during the shortest ongoing LGN ISIs and decreased as this value approached 30 ms (Figure 6B and C). This is to be expected given that most LGN spikes are triggered by retinal EPSPs and it takes approximately 30 ms for the LGN membrane potential to return to baseline after these depolarizations occur (Usrey et al., 1998; Carandini et al., 2007). However, consistent with the de-inactivation of T-channels during longer LGN ISIs, there was an increase in retinal spike efficacy during ongoing LGN ISIs >50 ms that was maintained for the longest recorded values (> 300 ms; Figure 6B and C; white noise: β_ISI_ = 2.11 ±0.12, p < 10^-5^, dist. = binomial; drifting grating: β_ISI_ = 9.41±0.15, p < 10^-5^, dist. = binomial). Consistent with past reports, retinal ISI also had a strong influence on retinal spike efficacy, reflecting temporal summation of multiple retinal EPSPs in the thalamus (Usrey et al., 1998, Alitto et al., 2018; Carandini et al., 2007). To account for this effect, we calculated retinal ISI-spike efficacy for three categories of ongoing LGN ISIs: short (< 30 ms), medium (> 30 ms and < 100 ms), and long (> 100 ms). From this it is evident that the two factors, retinal ISI and ongoing LGN ISI, interact to influence retinal spike efficacy (Figure 6D and E).

Given that T-potentials can last for tens of milliseconds (Destexhe and Sejnowski, 2001), we next quantified the time course of retinal spike efficacy modulation following relatively prolonged LGN ISIs (Figure 7A). For this analysis, we plotted retinal spike efficacy as a function of time following the initiation of a geniculate burst. For each burst, time zero was set 4 ms after the cardinal spike, thus excluding the increase in retinal spike efficacy caused by the definition of a thalamic burst (i.e., at least two spikes within 4 ms). Given the influence of RGC ISI on retinal spike efficacy, we also calculated the "expected efficacy” as if retinal efficacy was not influenced by the preceding geniculate ISI, but was instead determined only by the retinal ISI (see Materials and Methods).

**Figure 7.**
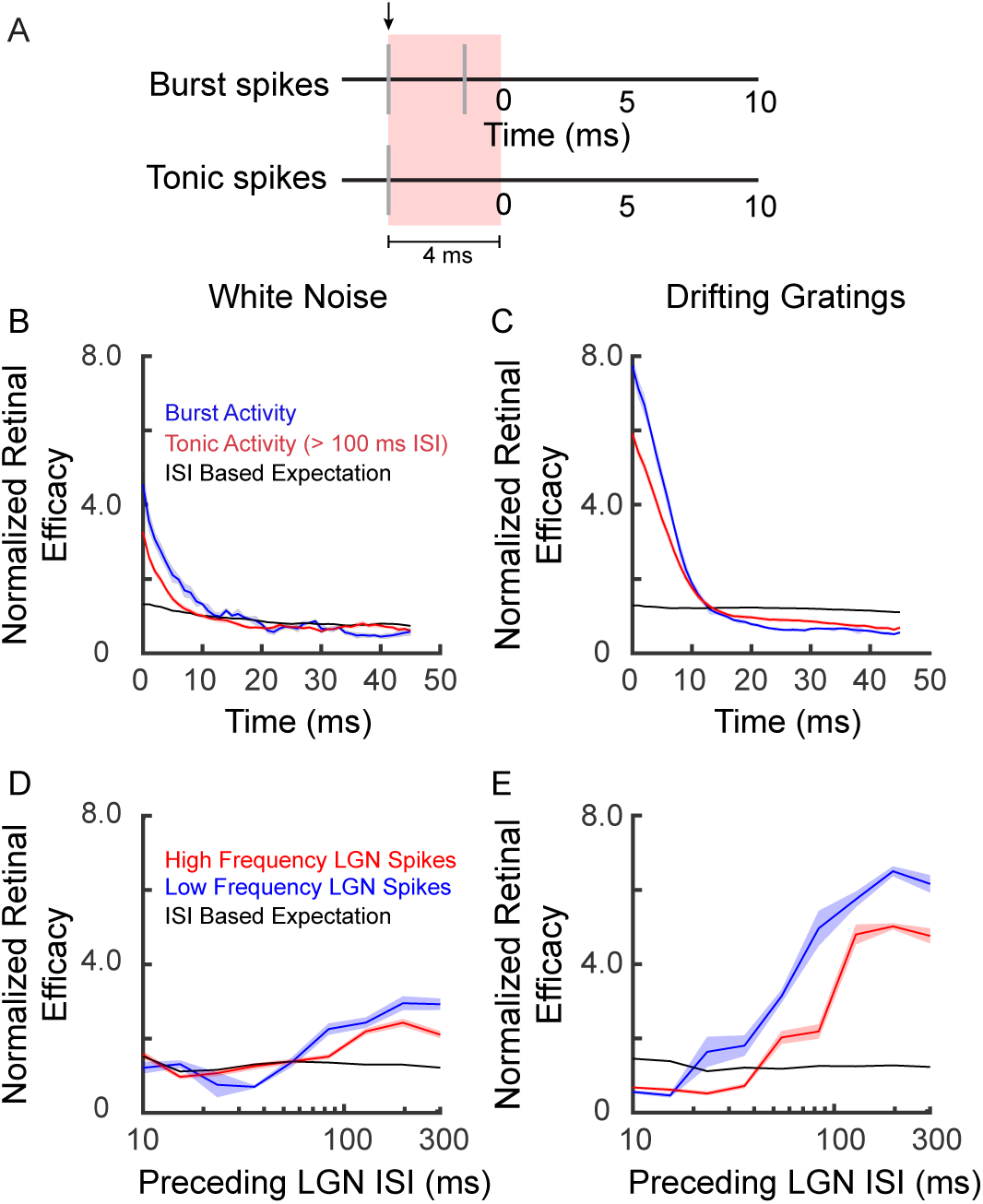
Retinal spike efficacy is influenced by preceding LGN ISI. ***A***, To quantify the influence of preceding LGN ISI on retinal spike efficacy, time = 0 was set to 4.0 ms after the cardinal spike in a burst or the referenced tonic spike (black arrow). *B*, *C*, Line plot showing that retinal spike efficacy is enhanced following both burst spikes (blue line) and tonic spikes with a preceding ISI > 100 ms (red line). The expected values given the preceding retinal ISIs are plotted as a baseline comparison (black line). Shaded areas indicate standard error. ***D***, ***E***, Line plots showing the influence of preceding ISI on retinal spike efficacy for 4 -10 ms following time 0 as indicated in ***A***.

Across our data set, there was a dramatic increase in retinal spike efficacy that lasted for approximately 10 ms from the onset of the burst compared to the expected efficacy values (Figure 7B and C; white noise: Burst β_0_ 0.55± = 0.34, Expected β_0_ -1.9± = 0.36, p < 10^-5^, dist. = binomial; drifting gratings: Burst β_0_ 2.1± = 0.40, Expected β_0_ -3.2± = 0.43, p < 10^-5^, dist. = binomial). Importantly, a similar modulation was seen for individual tonic spikes that were preceded by an ISI >100 ms (Figure 7B and C; white noise: Long IS Tonic β_0_ -3.2± = -0.11, p < 0.0005, dist. = binomial; drifting gratings: Long ISI Tonic β_0_ = 0.34 ± = 0.43, p < 10^-5^, dist. = binomial). Although, the increase in efficacy was greater following burst spikes compared to tonic spikes (white noise: p = 0.073; drifting gratings: p < 0.005), it is clear that the modulation of retinal efficacy is present even in the absence of classically defined bursts. As shown in Figure 7D and E the modulation of retinal spike efficacy is strongly dependent upon the LGN cells’ preceding ISI (white noise, high-frequency spikes: β_ISI_ 3.1±0.5, p < 10^-5^, low-frequency spikes β_ISI_ 3.4±1.8, p = 0.047; drifting gratings, high-frequency spikes: β_ISI_ 5.3±0.25, p < 10^-5^, low-frequency spikes β_ISI_ 4.0±0.2, p < 10^-5^). Further, the modulation of retinal efficacy begins to occur following geniculate ISIs shorter than would constitute a thalamic burst.

### Retinal contribution to geniculate burst spikes

Results presented above show that retinal spike efficacy is modulated by the preceding ISIs of LGN cells in a manner consistent with the involvement of T-channels and the amplification of visual signals within the LGN. To gain a comprehensive understanding of how response mode modulates retinogeniculate communication, we also quantified the influence of geniculate bursts on retinal contribution—the percentage of LGN spikes evoked from the recorded RGC. In general, geniculate bursts are expected to decrease retinal contribution by generating LGN spikes independent of retinal influence, therefore degrading the visual signal within the LGN. As described above, this is particularly true during geniculate bursts evoked by intrinsic corticothalamic oscillations. However, during visually driven geniculate bursts, such as the bursts examined in the current study (Figure 5), one may detect a decrease in retinal contribution, as measured via correlation analysis, even when there is no corresponding degradation of visual processing and LGN activity remains reliant on retinal influences. Thus, in addition to quantifying the influence of response mode on retinal contribution, we also sought to gain deeper insight into the functional consequences on visual processing in the LGN.

Consistent with the ability of T-channels to modulate retinal contribution, there was a significant inverse relationship between preceding LGN ISI and retinal contribution during visual stimulation (Figure 8A and B; white noise: β_ISI_ = -1.0±0.1, p = 10^-5^, dist. = binomial; drifting grating: β_ISI_ = -8.6±0.06, p < 10 ^-5^, dist. = binomial). This correlation was present even in the absence of high-frequency geniculate spikes (ISIs < 4 ms; white noise: β_ISI_ = -0.37±0.09, p = 0.0001, dist. = binomial; drifting grating: β_ISI_ = -4.4±1.2, p < 0.0001, dist. = binomial) and was evident for preceding ISIs < 100ms during visual stimulation with drifting gratings (drifting grating: β_ISI_ = -2.2±0.41, p < 10^-5^, dist. = binomial; white noise: β_ISI_ = -1.1±0.65, p =0.09, dist. = binomial), again reinforcing the conclusion that burst mode can influence geniculate activity even in the absence of classically defined bursts. For both white-noise and grating stimulation, the decrease in retinal contribution lasted for several milliseconds following a prolonged LGN ISI (Figure 8C and D, white noise = 6.1 ms; drifting gratings = 5.2 ms).

**Figure 8:**
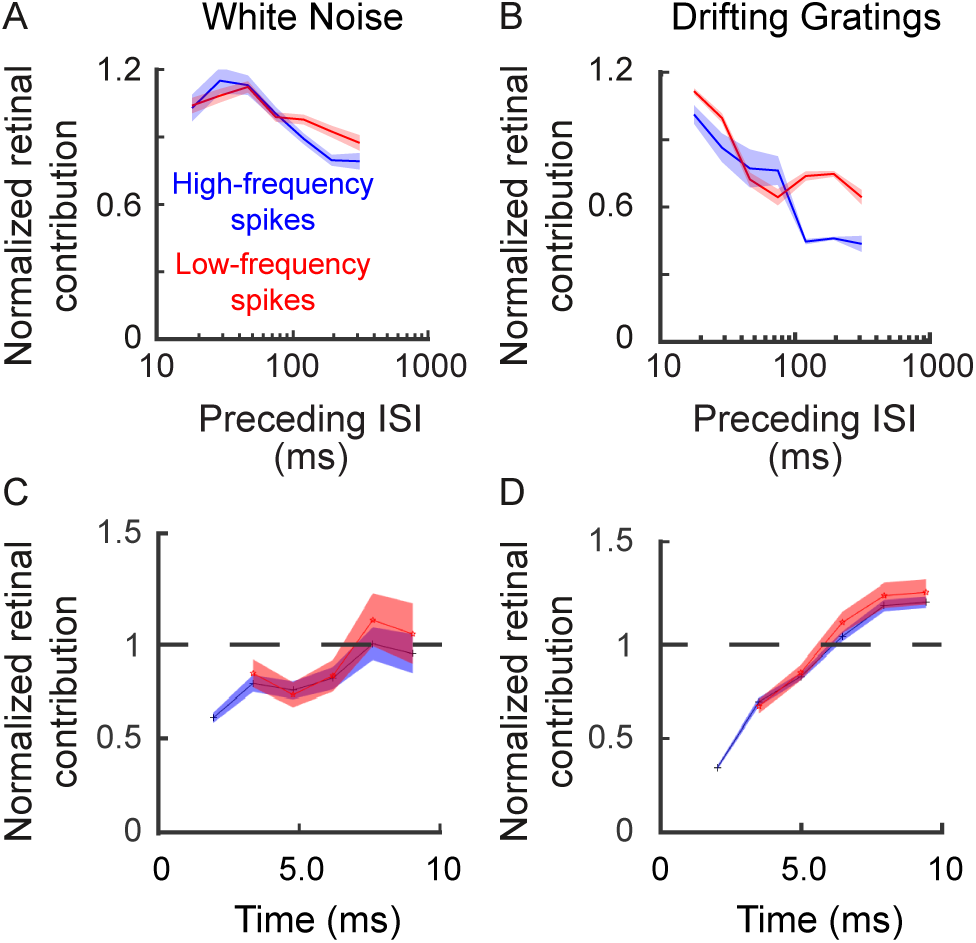
The influence of preceding ISI on retinal contribution. ***A***, ***B***, Line plot showing the influence of preceding LGN ISI on retinal spike contribution (red line = low frequency spikes, blue line = high frequency spikes). Shaded area indicates standard error. ***C***, ***D***, Line plots showing the temporal duration of the influence shown in ***A***, ***B***. Time zero is set as the occurrence of the initial spike following the referenced ISI (e.g., time of the cardinal spike in a burst).

We next wanted to determine if the measured decease in retinal contribution was caused by an uncoupling of retinal and geniculate activity. Although we measured a decrease in retinal contribution, it is possible that T-potentials allow single retinal spikes to evoke multiple LGN spikes, leading to an amplification of the retinal signal within the LGN. This would cause a decrease in the measured retinal contribution because the time delay from the triggering retinal spike increases with each subsequent LGN spike. Therefore, only the cardinal geniculate spike would fall into the monosynaptic window and thus be counted as triggered by the retina (Figure 9A). To determine the extent to which this occurred, we calculated retinal augmentation: here defined as the average retinal contribution minus the retinal contribution given that the previous spike was directly evoked by the recorded RGC. Effectively, this quantifies the relative change in contribution following an evoked spike. Positive values of retinal augmentation would be consistent with single retinal spikes triggering multiple LGN action potentials. Further, for retinal augmentation to be consistent with the involvement of T-potentials, then it should (1) increase with the preceding LGN ISI and (2) only be present during epochs containing relatively short subsequent LGN ISIs (e.g., high-frequency LGN spikes).

**Figure 9.**
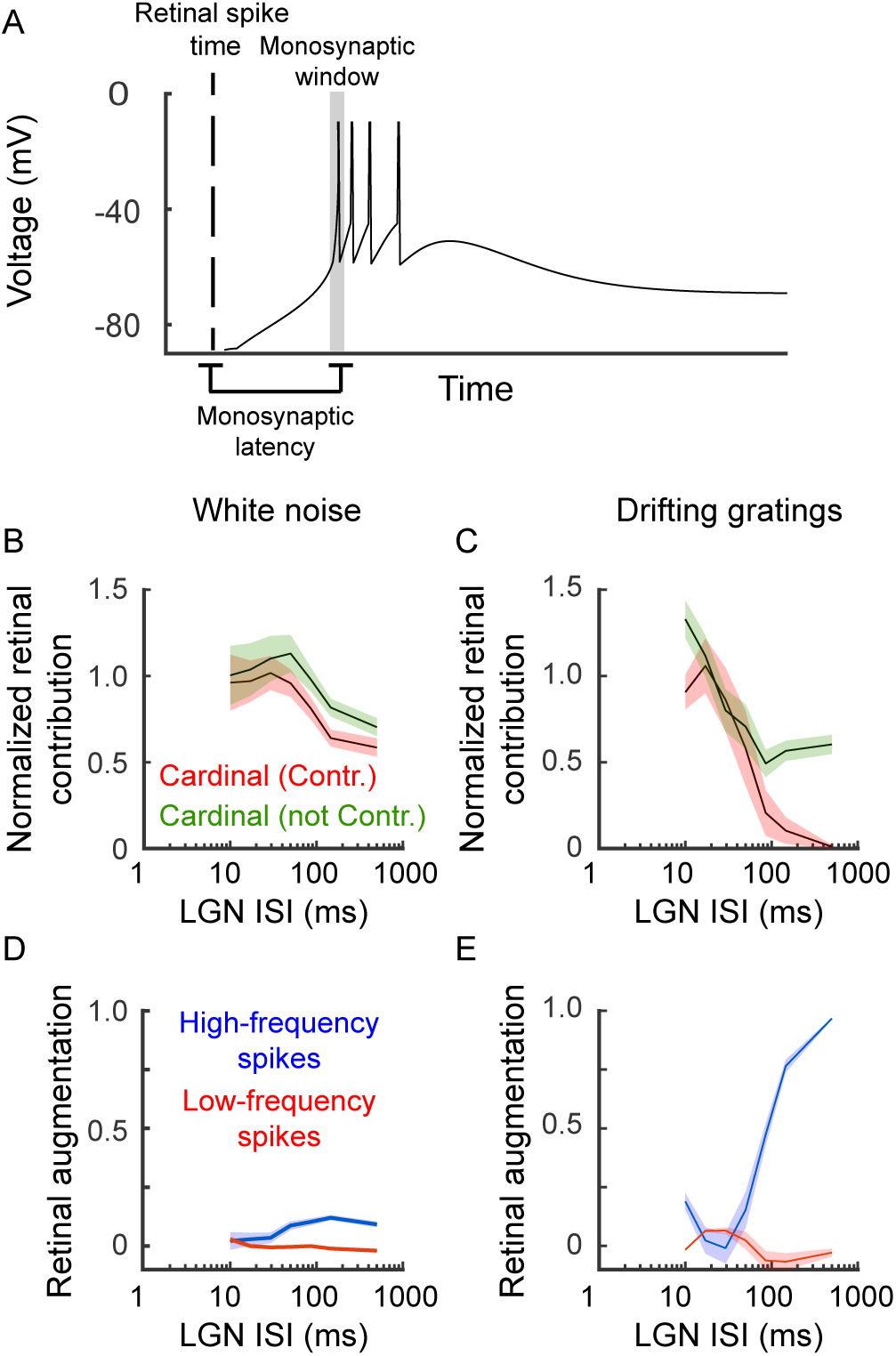
Augmentation of retinal transmission during high-frequency LGN activity. ***A***, When a retinal spike triggers a T-potential, it may result in a thalamic burst. Using correlation analysis only the cardinal spike would fall within the monosynaptic window (shaded box) and therefore be counted as driven by the retina. ***B***, ***C***, The influence of preceding LGN ISI on retinal contribution when the data is separated into two categories: cardinal spike was contributed by the recorded RGC (red line), cardinal spike was not contributed by the recorded RGC (green). Shaded area indicates standard error. ***D***, ***E***, Line plot showing retinal augmentation calculated from the data shown in ***B***, ***C***.

Retinal augmentation was significantly greater than zero during LGN bursts (Figure 9, blue trace, white noise: retinal augmentation = 0.23 ± 0.1, p = 0.0027; drifting gratings: retinal augmentation = 0.77 ± 0.13, p < 0.0001). Further, this effect was dependent upon the preceding LGN ISI, as measured by the difference in the influence of LGN ISI when the retinal contribution of the cardinal spike is considered (white noise: Cardinal Contributed, β_ISI_ = -2.8±0.34, Cardinal Not Contributed β_ISI_ = -1.7±0.40, p = 0.017, dist. = binomial: Cardinal Contributed β_ISI_ = - 24.0±0.81, Cardinal Not Contributed β_ISI_ = -3.6±0.16, p < 10^-5^, dist. = binomial). By comparison, in the absence of high-frequency LGN spikes, there was no evidence of signal augmentation, regardless of the preceding ISI (Figure 9D and E, red lines; white noise: retinal augmentation = 0.05 ± 0.06, p = 0.18; drifting gratings: retinal augmentation = -0.11 ± 0.12, p = 0.3).

The above analysis recasts the calculated decrease in retinal contribution for secondary burst spikes as an augmentation of the retinal signal; however, it does not address the decrease in retinal contribution for cardinal LGN burst spikes. One possible explanation for the decrease in retinal contribution for cardinal burst spikes is that T-potentials can be triggered by the release of inhibition, and this relationship is difficult to measure using correlation analysis. Burst spikes generated by the withdrawal of inhibition would lack a triggering retinal EPSP; however, they would still relay visual information to the cortex. To determine if LGN spikes that lacked a detectable triggering retinal spike encoded visual information, we calculated spike-count matched response maps for four categories of LGN spikes: contributed and non-contributed spikes during both tonic and burst response modes (Figure 10). While there was an overall decrease in signal-to-noise ratios for non-contributed spikes compared to contributed spikes (all contributed STN: 5.14± 0.54; all non-contributed STN: 3.5±0.41, p = 0.02), the decrease was present for both tonic spikes and burst spikes (non-contributed burst STN: 4.2± 0.6, non-contributed tonic STN: 2.8± 0.53, p = 0.1). Thus, there is no evidence that non-contributed burst spikes degrade the visual signal present in the LGN spike train.

**Figure 10.**
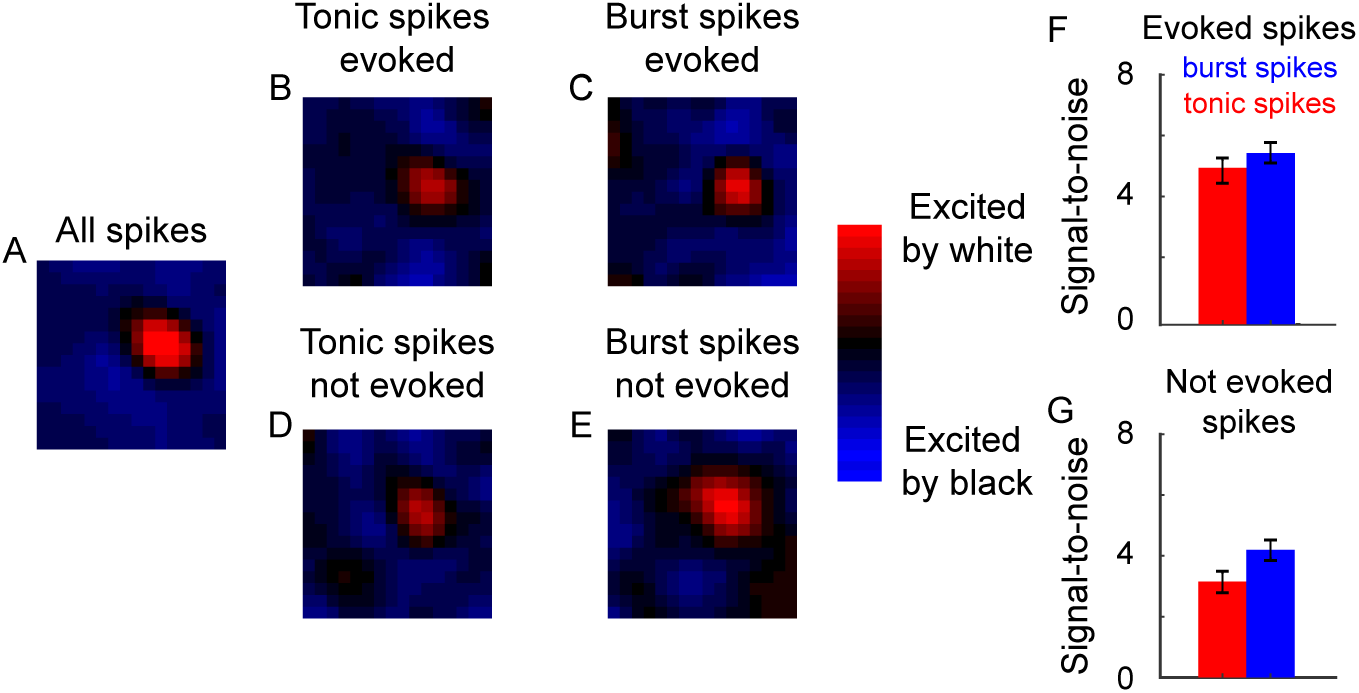
Burst spikes lacking a triggering RGC spike are nonetheless visually evoked. ***A***-***E***, STRFs calculated from different subsets of spike-count matched LGN spikes: (***A***) all spikes, (***B, C***) tonic and burst spikes evoked by the recorded RGC, (***D***, ***E***) tonic and bursts spikes that were not evoked by the recorded RGC. ***F***, ***G***, Bar graphs showing signal-to-noise ratios for LGN spikes that were either evoked (***F***) or not evoked (***G***)by the recorded RGC (red = tonic spikes, blue = burst spikes). Error bars indicated standard error.

## Discussion

The primary goal of this study was to determine the influence of thalamic burst mode on retinogeniculate communication. By simultaneously recording the spiking activity of monosynaptically connected pairs of RGCs and LGN neurons, we have shown that retinal signals to the cortex are amplified by visually evoked bursts in the LGN. This amplification is the result of (i) an increase in the probability that a retinal spike will trigger a geniculate response and (ii) an increase in the number of geniculate spikes that a single retinal spike can trigger. Further analysis demonstrates that the modulation of retinogeniculate communication increases as the preceding LGN ISI increases and the amplification of retinal activity occurs even in the absence of traditionally defined thalamic bursts. These results are consistent with the known biophysical properties of the T-type Ca^2+^ channels which underlie bursts in all thalamic nuclei (Llinás and Jahnsen,1982; Babadi, 2005; Destexhe, and Sejnowski, 2001; Sherman and Guillery, 2009; Elijah et al., 2015; Zeldenrust et al., 2018). We propose that T-potentials amplify the transmission of visual signals to primary visual cortex, most likely during periods of low-arousal. Given that this modulation can occur in the absence of thalamic bursts, T-potentials may also modulate retinogeniculate communication during behavioral conditions not typically associated with thalamic bursts.

### Retinogeniculate communication during visually driven LGN bursts can be explained by the known properties of T-type Ca^2+^ channels

Thalamic bursts are generated by the de-inactivation and subsequent activation of T-type Ca^2+^ channels (reviewed in Destexhe, and Sejnowski, 2001). This occurs when a strong hyperpolarization (de-inactivation) is followed by relatively rapid depolarization (activation). The depolarization can be active, as occurs with synaptic excitation, or passive, as occurs during the withdrawal of synaptic inhibition, or a combination of both (Andersen and Eccles, 1962; Llinás and Jahnsen, 1982; Hirsch et al., 1983; Deschenes et al., 1984; Destexhe, and Sejnowski, 2001). Importantly, the resulting Ca^2+^ mediated depolarizations (T-potentials) are dependent upon the depth and duration of the preceding hyperpolarization (Figure 3). Because of this, we hypothesized that T-channel activity could be estimated from the preceding LGN ISI. Consistent with this hypothesis, we found an increase in the probability and duration of high-frequency spiking (ISIs < 4 ms; the second criterion of a burst) as the preceding LGN ISI increased in duration (Figure 4). Likewise, there was a strong relationship between preceding LGN ISI and the amplification of retinogeniculate transmission (Figures 6-9).

During tonic response mode, it is generally assumed that each geniculate spike is triggered by a specific retinal action potential (Kaplan and Shapley, 1984; Sincich et al., 2007). As is common for monosynaptic interactions, and is particularly true at retinogeniculate synapses, cross-correlation analysis indicates that there is a precise monosynaptic window in which the triggering presynaptic spike can be found prior to the postsynaptic response (Figure 2). Although previous work has shown that the latency and duration of the monosynaptic window is invariant to changes in the visual stimulus (Fischer et al., 2017; Alitto et al., 2018), T-potentials can alter this relationship in two important ways. First, T-potentials can cause single retinal EPSPs to trigger multiple LGN action potentials. In this case, there is a retinal EPSP in the monosynaptic window of the cardinal burst spike; however, for each subsequent LGN spike, the triggering retinal spike occurs outside of this window (see Figure 9). Importantly, these burst spikes are all associated with retinal activity and should not be confused with intrinsically driven burst spikes that do not convey visual information. Second, T-potentials can be triggered by the release of inhibition, even in the absence of active depolarization (e.g., a retinal EPSP;). This is commonly referred to as a rebound potential. During visual stimulation, the withdrawal of inhibition often occurs with the onset of excitation (e.g., push-pull interactions; Wang et al., 2011; Suresh et al., 2016); however, the interaction of these two mechanisms may significantly transform the temporal relationship between retinal input and geniculate output, making it appear as if there was no triggering retinal excitation. As illustrated in Figure 10, burst spikes that did not have a detectable triggering retinal spike nonetheless convey visual stimulation and should not be confused with intrinsically driven activity that does not.

### Stimulus dependent amplification of visual signals

The biophysical properties of T-channels also explain stimulus-dependent differences in retinogeniculate communication during LGN bursts. In particular, the inferred influence of T-channels was greater with drifting grating stimulation compared to white-noise stimulation. This difference likely reflects the different spatiotemporal properties of drifting gratings and white-noise. Namely, the periodic nature of drifting gratings ensures that LGN neurons alternate between periods of strong excitation and strong inhibition, a pattern that is well suited for T-channel activity (Lu et al., 1992; Smith et al., 2000). By contrast, white-noise stimuli lack these correlations, leading to the generation of fewer and weaker T-potentials. Consistent with these differences, the amplification of visual signals was weaker and required longer geniculate ISIs during white-noise stimulation compared to drifting grating stimulation.

These stimulus dependent effects can also align our results with those from a previous study in macaque monkeys (Sincich et al., 2007). In this study, results indicate that retinal activity, as assessed using thalamic S-potentials, drives nearly all geniculate burst spikes. Similar to the white-noise stimulus used in the current study, the pink-noise stimulus used in the earlier study lacked the low-temporal frequencies that strongly de-inactivate T-channels, likely resulting in subthreshold T-potentials that were more dependent upon retinal excitation to drive geniculate action potentials. Although they did not examine the influence of geniculate bursts on retinal spike efficacy, if our suggestion is correct, then similar increases in retinal spike efficacy should be present in their data set. This would also indicate a shared mechanism across species to augment retinal signaling during geniculate bursts.

### Thalamic Burst Mode and Behavioral State

Visually evoked geniculate bursts are more likely to occur with inattentiveness or light anesthesia when the membrane potential of geniculate cells is thought to be more hyperpolarized than in the alert state. Under these conditions, the type of visual signal that is most likely to trigger a T-potential is a strongly suppressive stimulus followed by a strongly excitatory stimulus (Alitto et al., 2005). Resulting bursts effectively amplify the geniculocortical transmission of retinal signals resulting from the onset of a neuron’s preferred stimulus (Guido et al., 1992; Sherman and Guillery, 2002). In contrast, sleep and deep anesthesia engage intrinsic corticothalamic oscillations that dominate geniculate activity and drive synchronous bursting activity that serves to de-couple the thalamus and cortex from sensory activity (Steriade and Contreras, 1995; Timofeev et al., 1996; Elton 1997; Steriade, 2003). Thus, depending on the state of the corticothalamic circuitry, bursts may serve very different purposes: they can amplify the communication of visual signals to cortex during inattentiveness or light anesthesia or de-couple the thalamus and cortex during sleep and deep anesthesia.

Although bursts occur across all behavioral states, they occur most frequently during periods associated with diminished visual processing (Livingston and Hubel,1981; Bezdudnaya et al., 2006; Neil et al., 2010). With this in mind, the amplification of retinal signals during geniculate bursts should not be taken as evidence that visual processing is enhanced during periods of low arousal relative to periods of more highly engaged sensory processing. Rather, T-potentials enhance the ability of retinal spikes to trigger LGN activity during periods of otherwise diminished visual processing. During comparable behavioral states, a retinal EPSP that occurs during a T-potential is more likely to trigger LGN spikes than the same retinal spike in isolation. Given the relative suppression of tonic LGN activity during periods associated with geniculate bursts, the burst related retinal amplification functions as a contingency mechanism for the successful transmission of sensory signals to the cortex that would otherwise be lost.

Burst and tonic response modes are often described as binary states, which is an accurate description for the extreme ends of behavioral arousal: tonic mode during active sensory processing and burst mode during sleep and anesthesia. This hard distinction, however, fails to capture thalamic processing during the transition between the two response modes (Deleuze et al., 2012; Hong et al., 2014). In between the extremes of focused sensory processing and slow-wave sleep, the graded de-inactivation of a cell’s T-channels may play a previously underappreciated role in visual processing. Under certain conditions, the transition between tonic and burst response modes may approach a step function (Bezdudnaya et al. 2006); however, more studies are required to understand the full dynamic range of state dependent sensory processing. Finally, although bursts defined by classical criteria are less frequent in alert animals (Weyand et al., 2001; Ruiz et al., 2006; Weyand 2007; Alitto et al., 2011), this does not exclude the influence of T-potentials on visual responses in the LGN. T-potentials that do not trigger classically defined thalamic bursts may make a significant contribution to sensory processing in the engaged state.Alitto HJ, Moore BD 4th, Rathbun DL, Usrey WM (2011) A comparison of visual responses in the lateral geniculate nucleus of alert and anaesthetized macaque monkeys. J Physiol 589:87-99.

## Acknowledgements

We thank K.E. Neverkovec, D.J. Sperka, J. Johnson, and R. Oates for expert technical assistance. Supported by NIH grants EY013588 (WMU), P30 EY12576 (WMU), and the German Ministry for Education and Research, BMBF 031a308 (DLR).

## References

Alitto HJ, Rathbun DL, Fisher TG, Alexander PC, Usrey WM (2018) Contrast gain control and retinogeniculate communication. European Journal of Neuroscience. [Epub ahead of print; Mar 8, 2018; doi: 10.1111/ejn.13904]

Alitto HJ, Weyand TG, Usrey WM (2005) Distinct properties of stimulus-evoked bursts in the lateral geniculate nucleus. J Neurosci 25: 514–23.

Andersen P, Eccles J (1962) Inhibitory phasing of neuronal discharge. Nature 196: 645–647.

Babadi B (2005) Bursting as an effective relay mode in a minimal thalamic model. J Comput Neurosci 18: 229–243.

Bereshpolova Y, Stoelzel CR, Zhuang J, Amitai Y, Alonso JM, Swadlow HA (2011) Getting drowsy? Alert/nonalert transitions and visual thalamocortical network dynamics. J Neurosci 31: 17480–17487

Bezdudnaya T, Cano M, Bereshpolova Y, Stoelzel CR, Alonso JM, Swadlow HA (2006) Thalamic burst mode and inattention in the awake LGNd. Neuron 49: 421–32.

Carandini M, Horton JC, Sincich LC (2007) Thalamic Filtering of Retinal Spike Trains by Postsynaptic Summation. J Vision 7:20.1–11.

Cleland BG, Dubin MW, Levick WR (1971) Simultaneous recording of input and output of lateral geniculate neurones. Nat New Biol 231: 191–192.

Dan Y, Atick JJ, Reid RC (1996) Efficient coding of natural scenes in the lateral geniculate nucleus: experimental test of a computational theory. J Neurosci. 16:3351–3362.

Deleuze C, David F, Behuret, S, Sadoc G, Shin HS, Uebele V N, Renger JJ, Lambert RC, Leresche N, Bal T (2012) T-type calcium channels consolidate tonic action potential output of thalamic neurons to neocortex. J Neurosci 32: 12228–12236.

Denning KS, Reinagel P (2005) Visual control of burst priming in the anesthetized lateral geniculate nucleus. J Neurosci 25: 3531–8.

Deschenes, M., Paradis, M., Roy, J. P, Steriade, M (1984) Electrophysiology of neurons of lateral thalamic nuclei in cat: resting properties and burst discharges. J Neurophysiol 51: 1196–1219.

Destexhe A, Sejnowski, TJ (2001) Thalamocortical Assemblies. New York, NY: Oxford Press.

Destexhe A, Sejnowski, TJ (2002) The initiation of bursts in thalamic neurons and the cortical control of thalamic sensitivity. Philos Trans R Soc Lond B Biol Sci, 357: 1649–1657.

Elijah DH, Samengo I, Montemurro MA (2015) Thalamic neuron models encode stimulus information by burst-size modulation. Front Comput Neurosci, 9: 113.

Elton M, Winter O, Heslenfeld D, Loewy D, Campbell K Kok A (1997) Event-related potentials to tones in the absence and presence of sleep spindles. J Sleep Res 6: 78–83.

Guido W, Lu SM, Sherman SM (1992) Relative contributions of burst and tonic responses to the receptive field properties of lateral geniculate neurons in the cat. J Neurophysiol 68: 2199–2211.

Guido W, Lu SM, Vaughan JW, Godwin DW, Sherman SM (1995) Receiver operating characteristic (ROC) analysis of neurons in the cat’s lateral geniculate nucleus during tonic and burst response mode. Vis Neurosci 12: 723–741.

Hirsch, JC, Fourment, A, Marc ME (1983) Sleep-related variations of membrane potential in the lateral geniculate body relay neurons of the cat. Brain Res 259: 308–312.

Hong SZ., Kim HR, Fiorillo, CD (2014) T-type calcium channels promote predictive homeostasis of input-output relations in thalamocortical neurons of lateral geniculate nucleus. Front Comput Neurosci 8: 98.

Huguenard JR, McCormick DA (1992) Simulation of the currents involved in rhythmic oscillations in thalamic relay neurons. J Neurophysiol 68: 1373–1383.

Kaplan E, Shapley R (1984) The origin of the S (slow) potential in the mammalian lateral geniculate nucleus. Exp Brain Res 55: 111–116.

Lesica NA, Stanley GB (2004) Encoding of natural scene movies by tonic and burst spikes in the lateral geniculate nucleus. J Neurosci 24: 10731–10740.

Livingstone MS, Hubel DH (1981) Effects of sleep and arousal on the processing of visual information in the cat. Nature 291: 554–61.

Llinás R, Jahnsen H (1982) Electrophysiology of mammalian thalamic neurones in vitro. Nature 297: 406–8.

Lu SM, Guido W, Sherman SM (1992) Effects of membrane voltage on receptive field properties of lateral geniculate neurons in the cat: contributions of the low-threshold Ca2+ conductance. J Neurophysiol 68: 1285–1298.

Martinez LM, Molano-Mazón M, Wang X, Sommer FT, Hirsch JA (2014) Statistical wiring of thalamic receptive fields optimizes spatial sampling of the retinal image. Neuron 81: 943–56.

Niell CM, Stryker MP (2010) Modulation of visual responses by behavioral state in mouse visual cortex. Neuron 65: 472–479.

Ortuño T, Grieve KL, Cao R, Cudeiro J, Rivadulla C. 2014. Bursting thalamic responses in awake monkey contribute to visual detection and are modulated by corticofugal feedback. Front Behav Neurosci 8: 198.

Rathbun DL, Alitto HJ, Warland DK, Usrey WM (2016) Stimulus Contrast and Retinogeniculate Signal Processing. Front Neural Circuits 10: 8.

Rathbun DL, Warland DK, Usrey WM (2010) Spike timing and information transmission at retinogeniculate synapses. J Neurosci 30: 13558–13566.

Raudenbush SW, Bryk AS (2002) Hierarchical Linear Models: Applications and Data Analysis. London, England: Sage Publications.

Reid RC, Shapley RM (1992) Spatial structure of cone inputs to receptive fields in primate lateral geniculate nucleus. Nature 356: 716–718.

Reid RC, Victor JD Shapley RM (1997) The use of m-sequences in the analysis of visual neurons: linear receptive field properties. Vis Neurosci 16: 1015–1027.

Reinagel P, Godwin D, Sherman SM, Koch C (1999) Encoding of visual information by LGN bursts. J Neurophysiol 81, 2558–2569.

Ruiz O, Royal D, Sáry G, Chen X, Schall JD, Casagrande VA. 2006. Low-threshold Ca2+-associated bursts are rare events in the LGN of the awake behaving monkey. J Neurophysiol 95: 3401–3413.

Sherman SM, Guillery RW (2002) The role of the thalamus in the flow of information to the cortex. Philos Trans R Soc Lond B Biol Sci 357, 1695–1708.

Sherman SM, Guillery RW (2009) Exploring the Thalamus and its Role in Cortical Function, second edition. Cambridge, MA: MIT Press.

Sincich LC, Adams DL, Economides JR, Horton JC (2007) Transmission of Spike Trains at the Retinogeniculate Synapse. J Neurosci 27: 2683–2692.

Smith GD, Cox CL, Sherman SM, Rinzel J (2000) Fourier analysis of sinusoidally driven thalamocortical relay neurons and a minimal integrate-and-fire-or-burst model. J Neurophysiol 83: 588–610.

Steriade M (2003) The corticothalamic system in sleep. Front Biosci 8: 878–899.

Steriade M, Contreras D (1995) Relations between cortical and thalamic cellular events during transition from sleep patterns to paroxysmal activity. J Neurosci 15: 623–642.

Suresh V, Ciftcioglu UM, Wang X, Lala BM, Ding, K. R, Smith, WA, Sommer FT, Hirsch JA (2016) Synaptic Contributions to Receptive Field Structure and Response Properties in the Rodent Lateral Geniculate Nucleus of the Thalamus. J Neurosci. 36: 10949–10963.

Sutter EE (1987) A practical non-stochastic approach to nonlinear time-domain analysis. In: Advanced Methods of Physiological Systems Modeling. Los Angeles: University of Southern California, vol. 1, p. 303–315.

Swadlow HA, Gusev AG (2001) The impact of 'bursting' thalamic impulses at a neocortical synapse. Nat Neurosci 4: 402–408.

Timofeev I, Contreras D, Steriade M (1996) Synaptic responsiveness of cortical and thalamic neurons during various phases of slow oscillation in cat. J Physiol (Lond) 494: 265–278.

Usrey WM, Alitto HJ (2015) Visual Functions of the Thalamus. Annu Rev Vis Sci 1: 351–371.

Usrey WM, Alonso JM, Reid RC (2000) Synaptic interactions between thalamic inputs to simple cells in cat visual cortex. J Neurosci 20: 5461–5467.

Usrey WM, Reppas JB, Reid RC (1998) Paired-spike interactions and synaptic efficacy of retinal inputs to the thalamus. Nature 395: 384–387.

Usrey WM, Reppas JB, Reid RC (1999) Specificity and strength of retinogeniculate connections. J Neurophysiol 82: 3527–3540.

Usrey WM, Sceniak MP, Chapman B (2003) Receptive fields and response properties of neurons in layer 4 of ferret visual cortex. J Neurophysiol 89: 1003–1015.

Wang, X, Sommer, FT, Hirsch, JA (2011) Inhibitory circuits for visual processing in thalamus. Curr Opin Neurobiol 21: 726–733.

Wang, XJ, Rinzel, J, Rogawski MA (1991) A model of the T-type calcium current and the low-threshold spike in thalamic neurons. J Neurophysiol 66: 839–850.

Wei H, Bonjean M, Petry HM, Sejnowski TJ, Bickford ME (2011) Thalamic burst firing propensity: a comparison of the dorsal lateral geniculate and pulvinar nuclei in the tree shrew. J Neurosci 31: 17287–17299.

Weyand TG, Boudreaux M, Guido W (2001) Burst and tonic response modes in thalamic neurons during sleep and wakefulness. J Neurophysiol 85: 1107–1118.

Weyand TG (2007) Retinogeniculate transmission in wakefulness. J Neurophysiol 98: 769–785.

Zeldenrust F, Chameau P, Wadman WJ (2018) Spike and burst coding in thalamocortical relay cells. PLoS Comput Biol 14:2:e1005960. doi: 10.1371/journal.pcbi.1005960

